# A model to analyze age-related differences in human-generated head-tail sequences

**DOI:** 10.1101/2023.08.07.552223

**Authors:** Sergio Baena-Mirabete, Rosario García-Viedma, Sara Fernández-Guinea, Pere Puig

**Affiliations:** Department of Mathematics, Autonomous University of Barcelona, Barcelona, Spain; Department of Psychology, University of Jaén, Jaén, Spain; Department of Experimental Psychology, Complutense University of Madrid, Madrid, Spain

## Abstract

The generation of random-like sequences is a common task for assessing high-level cognitive abilities, such as inhibition, sustained attention and working memory. In general, many studies have shown a detrimental effect of aging on pseudo-random productions. The performance of participants in random generation tasks has typically been assessed by measures of randomness such as, among others, entropy and algorithmic complexity that are calculated from the series of responses produced by the subject. We focus on analyzing the mental model of randomness that people implicitly use when producing random series. We propose a novel latent class model based on Markov chains that aims to classify individuals into homogeneous classes according to the way they generate head-tail series. Our results reveal that there are significant age-related differences in the way individuals produce random-like sequences. Specifically, the group of healthy adults implicitly uses a simpler mental mechanism, in terms of memory requirements, compared to the group of younger participants.

**Author summary:** It is well known that, in general, people deviate from randomness as they attempt to mentally generate head-tail sequences as randomly as possible. The extensive literature on this topic has shown that human-generated head-tail series tend to have more alternations than would be expected by chance. However, it seems unrealistic to suppose that all individuals generate sequences based on the same random mental model. We conducted an experiment in which 331 individuals were asked to mentally simulate a fair coin: 69 healthy older adults with an age ≥ 60 and 262 Biology students with an age between 18 and 20. We found that the way in which random sequences are generated varies between subjects. A similar approach could be used to analyze differences in random generation tasks between subjects with different disorders and healthy subjects.

## Introduction

The generation of ‘random’ human sequences is a common task in experimental psychology and neuropsychology. Participants in random number generation tasks are asked to generate random-looking sequences, such as those involving binary digits (heads-or-tails). These tasks are commonly used to assess higher level cognitive abilities such as inhibition, sustained attention and working memory. When subjects are asked to produce ‘random’ sequences, the mental mechanism involved is as follows [1–3]: the storage capacity of the working memory is limited, so each subject can only remember a few previous responses. Based on these last items active in the working memory, a putative response is generated. This putative response is evaluated in conjunction with the previous responses to determine whether or not it is ‘random’. The next response is produced or inhibited based on this evaluation. If the individual inhibits the production of the putative response, this loop is repeated again. If the new response is produced, it is appended to the sequence in the working memory, while the first one is removed. All these selection and control functions correspond exactly to the role that is assigned to the central executive system of working memory [4–8].

Random number generation tasks have been used to study different disorders, such as schizophrenia [9, 10], autism [11], depression [12] and neurodegenerative syndromes such as Parkinson’s [13, 14] and Alzheimer’s disease [15]. All these studies showed that the sequences generated by the brain-damaged patients were more stereotyped than those produced by their respective control group. Other studies have also examined the effect of age on random generation, finding differences between young and elderly subjects [16, 17]. In general, these studies revealed age-related decrements in random generation performance. Interestingly, Gauvrit et al. [18] showed that the developmental curve of the ability to produce random-like sequences is similar to that of most cognitive abilities, with a peak of performance around age 25 and a decline starting around 60.

The performance of participants in random generation tasks is usually assessed by measures of randomness such as entropy, among others, which are calculated from the series of responses produced by the subject [19–21]. In practice, several of these measures are needed to obtain an acceptable diagnosis of the quality of randomness and, for this reason, they have been subject to strong criticism [22]. Some researchers have recently proposed using an approximation to Kolmogorov-Chaitin (algorithmic) complexity as a measure of randomness for short strings [23, 24]. The basic idea behind algorithmic complexity is that a string is random (or complex) if it cannot be produced by a program much shorter in length than the string itself. Each of these measures summarizes the degree of randomness (or complexity) of a given sequence in a single number and allows for simple comparison of results between subjects. However, these measures alone do not provide information about the mental mechanism employed to generate the sequences. In contrast, other studies have focused on analyzing the mental model of randomness that people use implicitly while producing random-like series. In this sense, a Markov chain of a given order is a well-known probabilistic model that is usually used to describe human-generated series [26–28]. These models assume that subjects’ responses depend only on their previous short-term choices, which is consistent with the limited capacity of working memory [29, 30]. In line with this approach, Meyniel et al. [31] suggested that the brain acts as a quasi-optimal inference device that constantly attempts to infer the transition probability matrix among the stimuli it receives and uses these inferences to generate statistical expectations about future observations.

Here, we attempt to analyze age-related differences in the way ‘random’ head-tail sequences are mentally produced. For this purpose, we propose a novel latent class model based on Markov chains. Latent class models are commonly used in behavioral and social science research [32–35] and, more specifically, for classification purposes [36, 37]. In general, latent class models are used to uncover unobserved heterogeneity in a population and find groups of individuals who respond similarly to the measured variables. Our proposed model aims to classify individuals into homogeneous classes according to the way they generate head-tail series and allows us to identify differences between age groups.

This article is structured as follows: first, we explain the experiment conducted with two groups of subjects, one of young students and the other of healthy older adults. Secondly, we present the theoretical framework and, finally, we describe the results obtained, as well as the main conclusions and discussion.

## Materials and methods

### The data

We conducted an experiment in which 331 individuals were asked to mentally simulate a fair coin: 69 healthy older adults (24 males, 45 females) with an age ≥ 60 and 262 Biology students (97 males, 165 females) with an age between 18 and 20.

As for the group of young students, the experiment was carried out at the Autonomous University of Barcelona on different days and in groups of around 30 individuals. Each session started early in the day and took place in the same classroom.

With regard to the group of older adults, the experiment took place in the psychology laboratories of the University of Jaén and the Complutense University of Madrid. The mean age was 70.89 years and the mean number of years of schooling was 9.52 years. The following inclusion criteria were established: Blessed’s dementia scale (BDS) ≤ 4, Mini-Mental State Examination (MMSE) ≥ 31, Geriatric Depression Scale (GDS) ≤ 5. In addition, the following exclusion criteria were imposed: presence or history of a chronic mental illness, neurological or cerebrovascular disorders, presence of a systemic disease that affects cognition or significant deficiencies in hand mobility.

After a briefing on the experiment and a practice session beforehand, each individual was asked to mentally produce and record a series of 50 head-tail outcomes as randomly as possible using a Windows dialog box designed with an Excel Visual Basic for Applications (Fig 1). To do this, the subjects were repeatedly required to press the key containing ‘smiley face figure’ for ‘HEADS’ and the key containing ‘cross figure’ for ‘TAILS’. It is important to emphasize that the results were recorded in a hidden worksheet so that the individual could not see the previous choices. To facilitate the analysis, the head-tail responses were recoded as a dichotomous variable with numerical values 0 (‘tails’) and 1 (‘heads’).

**Fig 1.**
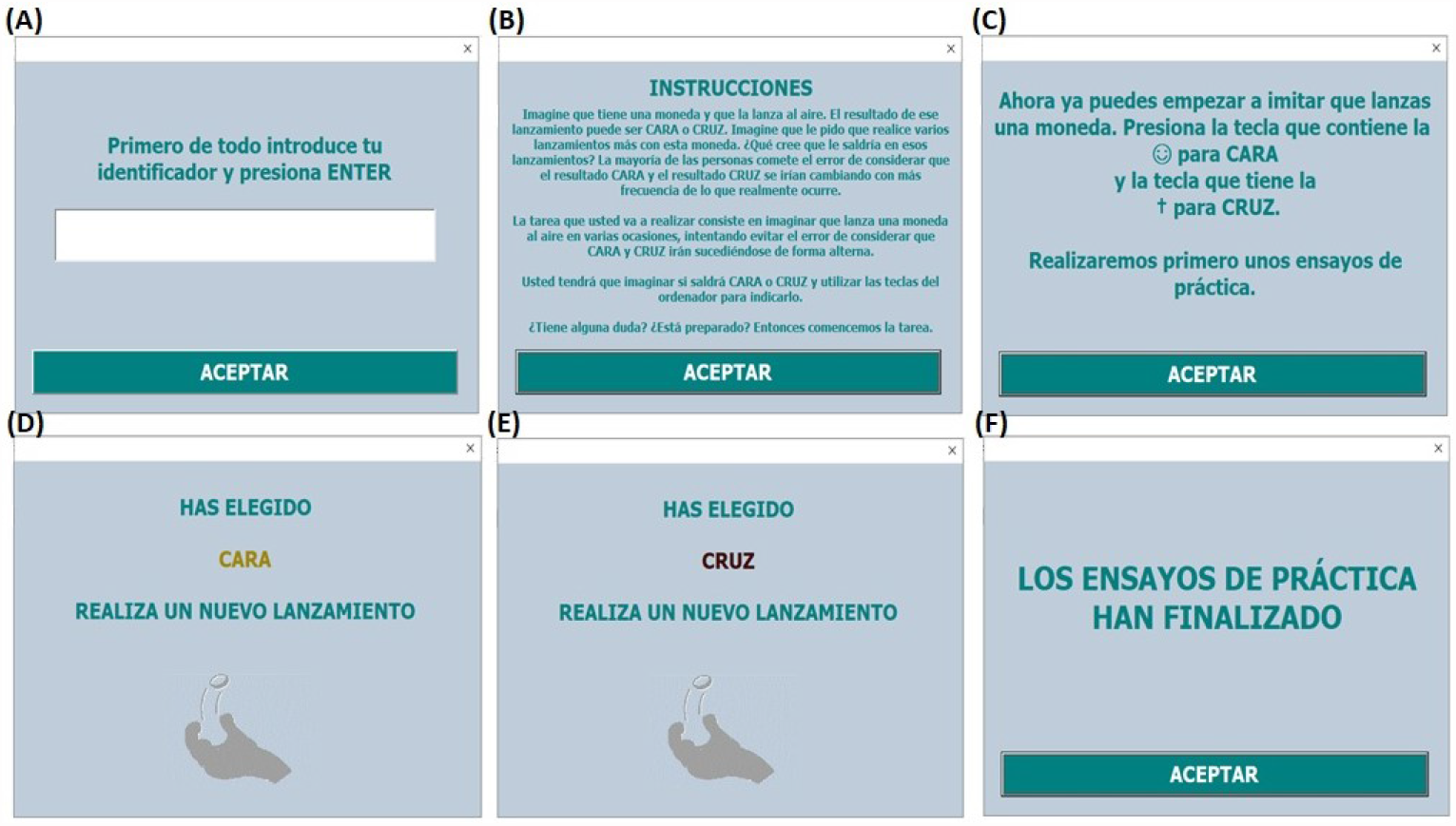
Screenshots of the interface seen by participants during the random number generation task. Panel A: “First, enter your identifier and press ENTER”. Panel B: “INSTRUCTIONS: Imagine you have a coin and you toss it in the air. The result of that toss can be either HEADS or TAILS. Imagine I ask you to make several more tosses with this coin. What do you think you would get on those tosses? Most people make the mistake of thinking that HEADS and TAILS alternate more often than they actually do by chance. The task you are about to perform is to imagine flipping a coin several times, trying to avoid the mistake of thinking that it will come up HEADS and TAILS alternately. Imagine whether the result will be HEADS or TAILS and use the keys on your computer to indicate this. Are you in doubt? Are you ready? Then let’s start the task”. Panel C: “You can now replicate the toss of a coin. Press the key containing ‘smiley face figure’ for HEADS and the key containing ‘cross figure’ for TAILS. First we will do some practice”. Panel D: “You have chosen HEAD. Make a new roll”. Panel E: “You have chosen TAIL. Make a new roll”. Panel F: “Practical tests have been completed”.

### Binary Markov chain model

Let {*X*_*t*_ : *t* = 1, …, *T* } be discrete-time random variables taking values in two possible states {0, 1}. Given that subscript *t* refers to a certain point in time, *X*_*t*_ is the corresponding state at time *t*, and *X*_*t*−*k*_ is the state *k* periods before (*k* < *t* ≤ *T*). Then, we have a *k*-order Markov chain if,

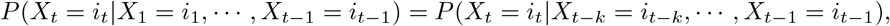

where *i*_1_, …, *i*_*T*_ ∈ {0, 1}. From now on, MC_*k*_ will denote a full parameterized *k*-order binary Markov chain model. Thus, MC_0_ corresponds to a series of independent outcomes. It is interesting to note that the full parameterized MC_*k*_ model can be described using a logistic model with

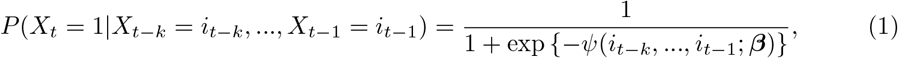

where ***β*** denotes the vector of parameters appearing in the linear predictor which has the form,

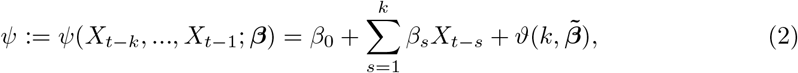

where 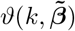 is the *k*-order function of interaction terms. Note that the number of parameters of these full parameterized MC_*k*_ models increases very rapidly as *k* grows: a binary MC_*k*_ model has 2^*k*^ parameters to be estimated [25].

Baena-Mirabete et al. [26] suggested a logistic autoregressive model for binary time series that takes into account interaction terms up to order *r* (*r* < *k*), as follows:

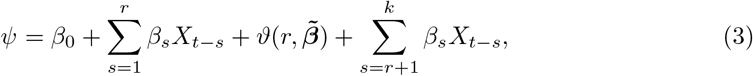

where 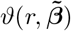 is the *r*-order function of the interaction terms and 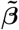 is the vector of parameters for these interaction terms. This model has 2^*r*^ + *k* − *r* parameters, fewer than the corresponding saturated MC_*k*_ model, which has 2^*k*^ parameters.

A more parsimonious model is the so-called *k-order short-long-time memory* Markov chain model of degree *r* (SLMC_*k*_(*r*)) [26], which can be obtained from Eq (3) under the hypothesis that *β*_*r*+1_ = … = *β*_*k*_ = *γ*. That is,

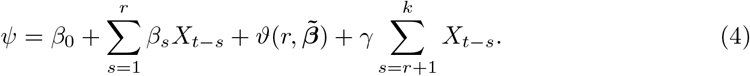

This model has 2^*r*^ + 1 parameters to be estimated. In the context of human-generated series, the SLMC_*k*_(*r*) model can be directly interpreted in the sense that it seems reasonable to believe that individuals are able to remember the immediately previous responses in an exact way, but only a summary of what happened many times before.

### Latent class model

Miller [29] argued that the number of elements an average human can hold in working memory is 7 ± 2. Subsequent studies corroborated this upper limit on our capacity to process information [30]. Consequently, in the analysis of human-generated head-tail series, it seems unrealistic to assume that individuals generate sequences based on a Markov process with equal memory-order for all of them.

Consider a sample of *N* individuals and, for the *j*th individual, the observed sequence of binary random variables over time **x**_**j**_ = {*X*_*jt*_ = *i*_*t*_ : *t* = 1, …, *T*}, *j* = 1, …, *N* and *i*_1_, …, *i*_*T*_ ∈ {0, 1}. We propose a binary Markov chain model whose parameters and memory-order differ across unobservable subgroups or latent classes. Let us denote the latent variable as *V*_*j*_, the number of latent classes as *C* and a particular class of the latent variable as *c*, with *c* = 1, 2, …, *C*. Let *k*_*c*_ be a positive integer such that 0 = *k*_1_ < *k*_2_ < … < *k*_*C*_. The latent class model proposed here considers that given a particular value of the latent variable *V*_*j*_ and the immediately *k*_*c*_ lagged outcomes, the observed binary variables over time are mutually independent. It is further assumed that each individual belongs to one of *C* exhaustive and mutually exclusive classes with probability *P* (*V*_*j*_ = *c*). The likelihood contribution, for the *j*th individual, is given by,

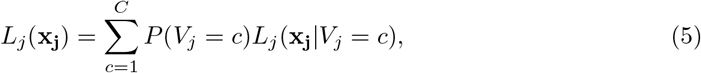

where *L*_*j*_ (**x**_**j**_|*V*_*j*_ = *c*) is the likelihood conditional on the latent class *c*. Specifically, we assume that *L*_*j*_ (**x**_**j**_|*V*_*j*_ = *c*) corresponds to the likelihood of a 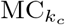. Note that for *c* = 1, we have the MC_0_ model that corresponds to a series of independent outcomes.

To simplify the subsequent notation, let us define the transition probabilities conditioned on the latent class *c* as

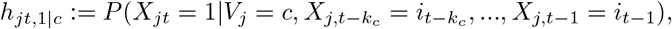

for *c* = 2, 3, …, *C*. Then, the Bernoulli distribution with probability of success *h*_*jt*,1|*c*_ is given by:

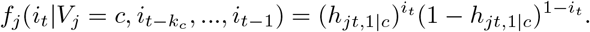

Similarly, the success probability conditioned on the latent class *c* = 1, corresponding to a series of independent outcomes, is defined as

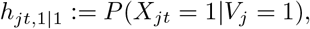

and *f*_*j*_ (*i*_*t*_|*V*_*j*_ = 1) denotes the Bernoulli distribution with constant probability *h*_*jt*,1|1_. We can now express the likelihood contribution, for the *j*th individual and *t* ≥ *k*_*C*_ + 1, from Eq (5) as follows:

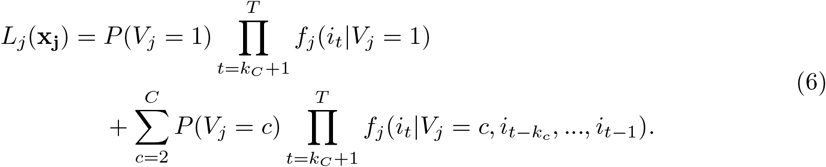

Since individuals are assumed to be independent, the global likelihood is given by:

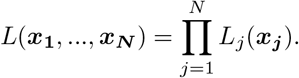

The parameters to estimate are the class membership probabilities *P* (*V*_*j*_ = *c*) and the probabilities *h*_*jt*,1|*c*_, *c* = 1, 2, …, *C*, which are stated using logistic functions (Eq (1)). The EM algorithm [38] has been used to obtain the maximum likelihood estimates for the parameters.

The probability of *j*th individual of belonging to a particular latent class *c* given the responses **x**_**j**_, i.e., the posterior membership probability, can be obtained by the Bayes rule as follows:

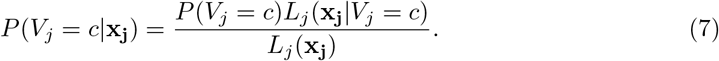

As a classification rule, we use the modal assignment, which amounts to assigning each individual to the latent class with the highest posterior probability, i.e,

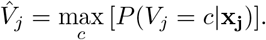

## Results

In this section, we present the results obtained by fitting the models described above to the sample of random-like series produced by the groups of young students and healthy older adults. First, we fit Markov chain models separately for both groups and obtain the main fit statistics. Second, we estimate latent class models based on Markov chains to uncover unobservable heterogeneity that may exist within groups and identify differences between both groups of age in the the way they generate head-tail series.

### Fitted Markov chain models: group of young students

Table 1 shows the goodness-of-fit statistics for various models based on Markov chains that are fitted to the sample of young students. According to the BIC-criterion, the best model is SLMC_6_(3) (Eqs (1) and (4)) which exactly takes into account the three previous head-tail choices, and a summary of the outcomes between the instants *t* − 4 and *t* − 6. In fact, short-memory models are not among those with the best fit statistics. In this regard, note, for example, that the SLMC_3_(1) model is nested in the SLMC_6_(3) model. The Likelihood Ratio Test found enough statistical evidence to reject the SLMC_3_(1) model in favour of the SLMC_6_(3) model (*p*-value <0.0001).

**Table 1.** BIC-statistics and the number of parameters, *q*, for the fitted Markov chains.

| Model | $q$ | $-\log L$ | BIC |
| --- | --- | --- | --- |
| MC<sub>0</sub> | 1 | 7809 | 15627 |
| MC<sub>1</sub> | 2 | 7766 | 15551 |
| MC<sub>2</sub> | 4 | 7711 | 15458 |
| MC<sub>3</sub> | 8 | 7670 | 15414 |
| MC<sub>4</sub> | 16 | 7606 | 15360 |
| MC<sub>5</sub> | 32 | 7538 | 15375 |
| SLMC<sub>2</sub>(1) | 3 | 7711 | 15449 |
| SLMC<sub>3</sub>(1) | 3 | 7710 | 15448 |
| SLMC<sub>4</sub>(1) | 3 | 7703 | 15433 |
| SLMC<sub>5</sub>(1) | 3 | 7679 | 15387 |
| SLMC<sub>6</sub>(1) | 3 | 7672 | 15372 |
| SLMC<sub>6</sub>(2) | 5 | 7657 | 15360 |
| SLMC<sub>6</sub>(3) | 9 | 7629 | 15342 |
| SLMC<sub>7</sub>(1) | 3 | 7677 | 15381 |
| SLMC<sub>7</sub>(2) | 5 | 7657 | 15360 |

$L$ is the estimated likelihood. The first seven observations for each subject are removed from the sample.

Parameter estimates are shown in Table 2. Note that all the estimated parameters are statistically significant (*p*-value <0.0001). The negative sign on the slope parameters (*X*_*t*−1_, *X*_*t*−2_ and *X*_*t*−3_) indicates that the probability of alternating responses is higher than the probability of persistent responses by producing, therefore, alternations from heads to tails (and vice versa) more often than really happen at random.

**Table 2.** Estimated parameters for the SLMC_6_(3) model.

| Parameter | Estimate | Standard error | $t$-statistic | $p$-value |
| --- | --- | --- | --- | --- |
| intercept | 1.108 | 0.078 | 14.278 | <0.0001 |
| $X_{t-1}$ | -0.771 | 0.082 | -9.441 | <0.0001 |
| $X_{t-2}$ | -0.835 | 0.082 | -10.174 | <0.0001 |
| $X_{t-3}$ | -0.569 | 0.082 | -6.969 | <0.0001 |
| $X_{t-1} \cdot X_{t-2}$ | 0.630 | 0.109 | 5.781 | <0.0001 |
| $X_{t-1} \cdot X_{t-3}$ | 0.556 | 0.109 | 5.111 | <0.0001 |
| $X_{t-2} \cdot X_{t-3}$ | 0.625 | 0.109 | 5.743 | <0.0001 |
| $X_{t-1} \cdot X_{t-2} \cdot X_{t-3}$ | -1.169 | 0.155 | -7.532 | <0.0001 |
| $\sum_{s=4}^6 X_{t-s}$ | -0.236 | 0.025 | -9.520 | <0.0001 |

The first six observations for each subject are removed from the sample.

Table 3 shows the probability of the occurrence of ‘head’ resulting from the estimated SLMC_6_(3) model given a particular sequence of previous head-tail responses. Interestingly, the higher the number of ‘heads’ in previous choices, the lower the estimated probability of ‘head’ in the current state. Take note, however, of the higher probability of ‘head’ obtained with the S_4_ sequence of preceding states than with the S_2_ sequence. Although both sequences have the same number of ‘heads’, they differ in the ordering of the three immediate head-tail choices. These results contrast with those that would be obtained with the independence model (MC_0_), which exactly mimics a fair coin toss with a 0.5 probability of turning up ‘heads’ regardless of previous outcomes.

**Table 3.**
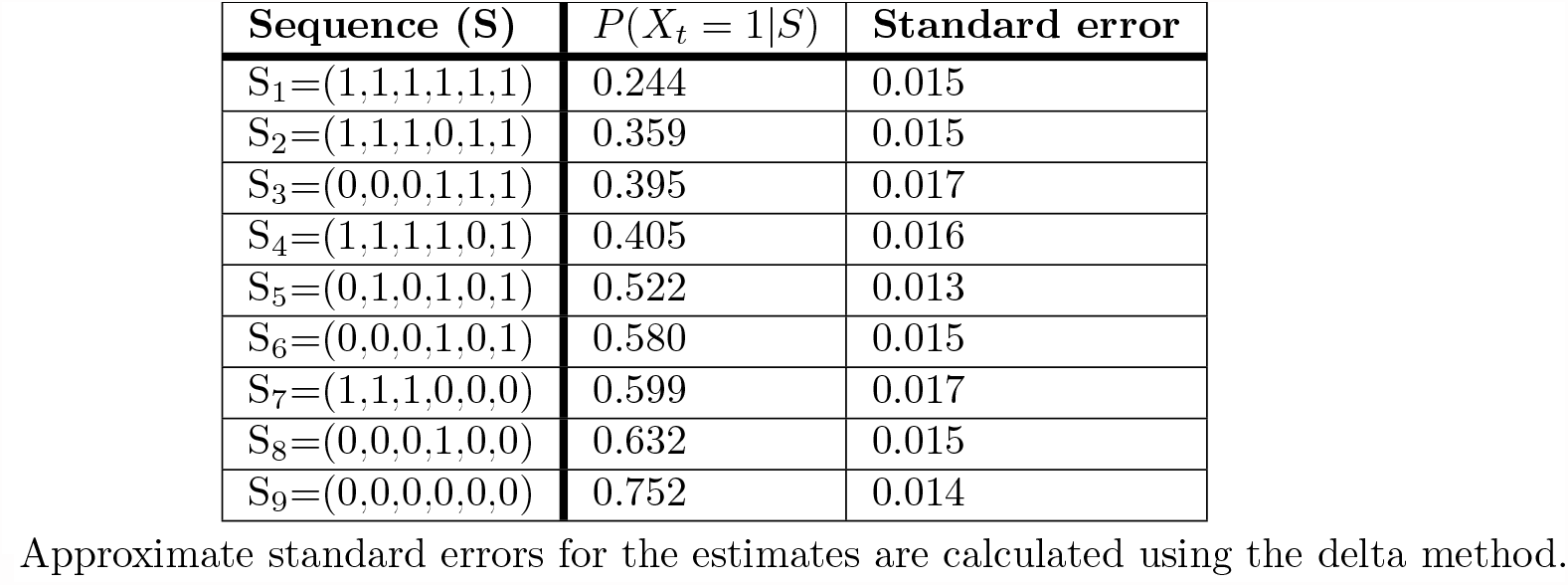
Probability of ‘head’ estimated by the SLMC_6_(3) model given the previous sequence of states *S* = (*i*_*t*−6_, *i*_*t*−5_, …, *i*_*t*−1_).

The SLMC_6_(3) model is a restricted version of the model defined in Eq (3) under the null hypothesis *H*_0_ : *β*_4_ = *β*_5_ = *β*_6_ = *γ*. The Likelihood Ratio Test did not find enough statistical evidence to reject it (*p*-value=0.180). These results suggest that the young students in the sample are generally able to remember in detail the sequence of their three immediately preceding choices, but only vaguely recall a summary of their longer-term responses up to memory-order 6.

### Fitted Markov chain models: group of healthy older adults

Table 4 shows the goodness-of-fit statistics for various Markov chain models that are fitted to the sample of healthy older adults. According to the BIC-criterion, the best model is SLMC_3_(1) (Eqs (1) and (4)) which exactly takes into account the immediately previous head-tail response, and a summary of the outcomes between the instants *t* − 2 and *t* − 3. Long-memory models, in contrast to the results obtained for the group of young students, are not among those with the best fit statistics. As mentioned previously, the SLMC_3_(1) model is nested in the SLMC_6_(3) model. The Likelihood Ratio Test did not find enough statistical evidence to reject the SLMC_3_(1) model in favour of the SLMC_6_(3) model (*p*-value=0.246).

**Table 4.** BIC-statistics and the number of parameters, *q*, for the fitted Markov chains.

| Model | q | -log L | BIC |
| --- | --- | --- | --- |
| MC<sub>0</sub> | 1 | 2104 | 4217 |
| MC<sub>1</sub> | 2 | 2089 | 4195 |
| MC<sub>2</sub> | 4 | 2079 | 4190 |
| MC<sub>3</sub> | 8 | 2054 | 4172 |
| MC<sub>4</sub> | 16 | 2031 | 4190 |
| SLMC<sub>2</sub>(1) | 3 | 2080 | 4183 |
| SLMC<sub>3</sub>(1) | 3 | 2062 | 4147 |
| SLMC<sub>4</sub>(1) | 3 | 2070 | 4163 |
| SLMC<sub>5</sub>(1) | 3 | 2070 | 4164 |
| SLMC<sub>6</sub>(1) | 3 | 2080 | 4184 |
| SLMC<sub>3</sub>(2) | 5 | 2061 | 4161 |
| SLMC<sub>4</sub>(2) | 5 | 2068 | 4176 |
| SLMC<sub>5</sub>(2) | 5 | 2067 | 4175 |
| SLMC<sub>6</sub>(2) | 5 | 2075 | 4190 |
| SLMC<sub>6</sub>(3) | 9 | 2054 | 4180 |

L is the estimated likelihood. The first six observations for each subject are removed from the sample.

Parameter estimates are shown in Table 5. Note that all the estimated parameters are statistically significant (*p*-value <0.0001). The negative sign on the slope parameters indicates that adult individuals deviate from randomness due to a greater tendency to alternate responses than would be the case in a random process.

**Table 5.**
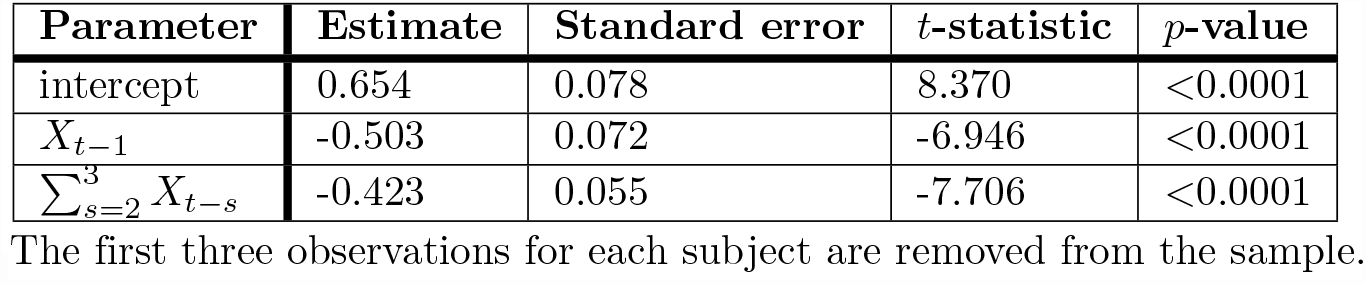
Estimated parameters for the SLMC_3_(1) model.

| Parameter | Estimate | Standard error | $t$-statistic | $p$-value |
| --- | --- | --- | --- | --- |
| intercept | 0.654 | 0.078 | 8.370 | $<0.0001$ |
| $X_{t-1}$ | -0.503 | 0.072 | -6.946 | $<0.0001$ |
| $\sum_{s=2}^3 X_{t-s}$ | -0.423 | 0.055 | -7.706 | $<0.0001$ |

The first three observations for each subject are removed from the sample.

Table 6 shows the probability of the occurrence of ‘head’ resulting from the estimated SLMC_3_(1) model given a particular sequence of previous head-tail responses. The results are comparable to those obtained in Table 3 for the same sequences. According to this model, healthy adults, in their mental process of generating a random series of heads-or-tails, only take into account, in their current decision, the last three choices. In this regard, it is worth noting that the estimated probabilities of ‘head’ given the sequences S_7_ and S_9_ are identical.

**Table 6.** Probability of ‘head’ estimated by the SLMC_3_(1) model given the previous sequence of states *S* = (*i*_*t*−6_, *i*_*t*−5_, …, *i*_*t*−1_).

| Sequence (S) | $P(X_t = 1\vert S)$ | Standard error |
| --- | --- | --- |
| S<sub>1</sub>=(1,1,1,1,1,1) | 0.333 | 0.018 |
| S<sub>2</sub>=(1,1,1,0,1,1) | 0.433 | 0.013 |
| S<sub>3</sub>=(0,0,0,1,1,1) | 0.333 | 0.018 |
| S<sub>4</sub>=(1,1,1,1,0,1) | 0.433 | 0.013 |
| S<sub>5</sub>=(0,1,0,1,0,1) | 0.433 | 0.013 |
| S<sub>6</sub>=(0,0,0,1,0,1) | 0.433 | 0.013 |
| S<sub>7</sub>=(1,1,1,0,0,0) | 0.658 | 0.018 |
| S<sub>8</sub>=(0,0,0,1,0,0) | 0.558 | 0.012 |
| S<sub>9</sub>=(0,0,0,0,0,0) | 0.658 | 0.018 |

Approximate standard errors for the estimates are calculated using the delta method.

The SLMC_3_(1) model is a restricted version of the model defined in Eq (3) under the null hypothesis *H*_0_ : *β*_2_ = *β*_3_ = *γ*. The Likelihood Ratio Test did not find enough statistical evidence to reject it (*p*-value=0.760). These results suggest that older adults, in their random generation task, only accurately remember their immediate choice and more vaguely their most distant responses up to memory-order 3.

### Fitted latent class models

As shown in the previous sections, the group of young students, in general, generate random-like sequences based on a random mental model of long-memory (order 6). In contrast, the group of healthy older adults rely on a random mental model of shorter memory (order 3). However, the approach presented earlier does not take into account the heterogeneity that may exist within the two groups. In the following, we assume that there can be 3 types of individuals according to their way of mentally generating random sequences: individuals ‘without memory’ who would be able to replicate a series of fair coin tosses (class 1), individuals of short-time memory (class 2) and of long-time memory (class 3). Table 7 shows the specification of a 3-latent class model whose general form of the likelihood function is detailed in Eq (6). Individuals belonging to class 2 will deviate from randomness since their current ‘heads-or-tails’ decision would be influenced by their immediate short-term choices. Individuals in class 3, in their random generation task, will show a deviation from randomness due to the influence not only of their immediate short-term choices, but also of their long-term responses, which they would at least vaguely be able to remember.

**Table 7.** Specification of a 3-latent class model of long-memory order *u* ≥4.

| Latent class | Class probabilities | Underlying process | Probability of ‘head’ ($h_{jt,1\vert c}$) |
| --- | --- | --- | --- |
| Class $c = 1$ | $P_{ad}(V_j = 1)$<br>$P_{st}(V_j = 1)$ | MC<sub>0</sub> | $h_{jt,1\vert 1} = \text{logit}(\beta_{01})$ |
| Class $c = 2$ | $P_{ad}(V_j = 2)$<br>$P_{st}(V_j = 2)$ | SLMC<sub>3</sub>(1) | $h_{jt,1\vert 2} = \text{logit}(\beta_{02} + \beta_{12}X_{j,t-1} + \beta_{22} \sum_{s=2}^3 X_{j,t-s})$ |
| Class $c = 3$ | $P_{ad}(V_j = 3)$<br>$P_{st}(V_j = 3)$ | SLMC<sub>$u$</sub>(1) | $h_{jt,1\vert 2} = \text{logit}(\beta_{03} + \beta_{13}X_{j,t-1} + \beta_{23} \sum_{s=2}^u X_{j,t-s})$ |

The subscripts of the class probabilities refer to the group (st=students, ad=older adults). The class probabilities sum to 1 for each of the groups. The probabilities of ‘head’ are modeled using a logistic link function: $\text{logit}(\psi) = 1/(1 + \exp\{-\psi\})$ .

Table 8 shows the goodness-of-fit statistics for several 3-latent class fitted models of different long-memory order (*u* ≥ 4). Based on the BIC-criterion, the best model is obtained for the long-memory order *u* = 6. Table 9 shows the estimates obtained for the model whose slope parameters are the same between the two groups and only the class probabilities differ. The Likelihood Ratio Test showed no strong evidence to reject the null hypothesis of equality of slope parameters between the two groups (*H*_0_ : ***β***_***st***_ **= *β***_***ad***_), with a *p*-value very close to the region of acceptance at the 95% confidence level (*p*-value = 0.048). Estimated slope parameters for each of the groups can be found in the Supporting Information (S1 Table and S2 Table). Again, the negative sign of the slope parameters in Table 9 indicates a greater tendency to produce sequences with more alternations than would occur by chance. Individuals who mentally generates head-tail sequences according to the underlying processes associated with latent classes 2 and 3 will produce an excess of alternations (negative sign in *b*_12_ and *b*_13_) and a tendency to balance the distribution of their responses in short sequence fragments (negative sign in *b*_22_ and *b*_23_). However, in this attempt to keep the distribution of their responses balanced and to evaluate randomness in fragments of short sequences, the mental mechanism implicit in the underlying process associated with latent class 3 is somewhat more sophisticated than that of class 2, since it uses a larger chunk of memory (up to memory-order 6) in which evaluations are performed. Finally, individuals who mentally generate a sequence according to the underlying process associated with latent class 1 will produce a series of not only independent but also balanced outcomes, as indicated by the near-zero value of the intercept (*b*_01_). According to the Likelihood Ratio Test, the class probabilities are statistically different between the groups of young students and healthy older adults (*p*-value < 0.0005). Interestingly, the probability of belonging to the latent class 2 is significantly higher in the group of healthy older adults compared to the group of young students.

**Table 8.** BIC-statistics and the number of parameters, *q*, for several fitted latent class models of different long-memory order (*u*).

| u | q | -log L | BIC |
| --- | --- | --- | --- |
| 4 | 11 | 9647 | 19398 |
| 5 | 11 | 9609 | 19323 |
| 6 | 11 | 9607 | 19319 |
| 7 | 11 | 9622 | 19349 |

L is the estimated likelihood. The first seven observations for each subject are removed from the sample.

**Table 9.** Estimated parameters for the latent class model of long-memory order *u* = 6.

| Parameter | Class 1 (c=1) | Class 2 (c=2) | Class 3 (c=3) |
| --- | --- | --- | --- |
| $P_{ad}(V_j = c)$ | 0.294 (0.078) | 0.312 (0.090) | 0.394 (0.106) |
| $P_{st}(V_j = c)$ | 0.366 (0.058) | 0.065 (0.026) | 0.569 (0.068) |
| $b_{01}$ | -0.029 (0.030) | | |
| $b_{02}$ | | 2.184 (0.287) | |
| $b_{12}$ | | -1.236 (0.194) | |
| $b_{22}$ | | -1.502 (0.192) | |
| $b_{03}$ | | | 1.770 (0.142) |
| $b_{13}$ | | | -0.901 (0.084) |
| $b_{23}$ | | | -0.549 (0.046) |

The first six observations for each subject are removed from the sample. Standard errors for the estimates in parentheses.

The number of individuals assigned to each latent class is shown in Table 10. Here we use the modal assignment that classifies each individual according to the highest posterior probability of belonging to each latent class (Eq (7)). Note that only 16 of the 262 young students (6.11%) were associated with a simpler mental process based on latent class 2, compared to the group of healthy adults, where the percentage was higher (30.43%). Most of the young students (61.45%), 161 out of 262 subjects, were assigned to latent class 3, related to a mental process with higher memory demands. Finally, a somewhat higher percentage of young students, compared to the healthy adult group, were assigned to latent class 1 (32.44% vs. 23.19%). The ternary diagrams (Fig 2) graphically represent the posterior probabilities of each individual belonging to each latent class as positions in an equilateral triangle. In a ternary diagram, a specific individual positioned at the triangle’s centroid has the same probability of belonging to each of the latent classes, i.e., 1/3. On the other hand, an individual located at one of the vertices of the triangle will have a probability equal to 1 of belonging to a given latent class and 0 in the other two classes. Therefore, a ternary plot in which individuals were located near the centroid of the triangle would indicate a very weak assignment of each individual to each of the latent classes. Conversely, a representation such as the one in Fig 2, in which subjects are distributed in the vicinity of the vertices of the triangle, indicates a strong assignment to each latent class.

**Table 10.** Number of individuals (and percentage) assigned to each latent class.

| | $\hat{V}_j = 1$ | $\hat{V}_j = 2$ | $\hat{V}_j = 3$ |
| --- | --- | --- | --- |
| No. of adults | 16 | 21 | 32 |
| % adults | 23.19 | 30.43 | 46.38 |
| No. of students | 85 | 16 | 161 |
| % students | 32.44 | 6.11 | 61.45 |

Each individual has been assigned to the latent class with the highest posterior probability (modal assignment).

**Fig 2.**
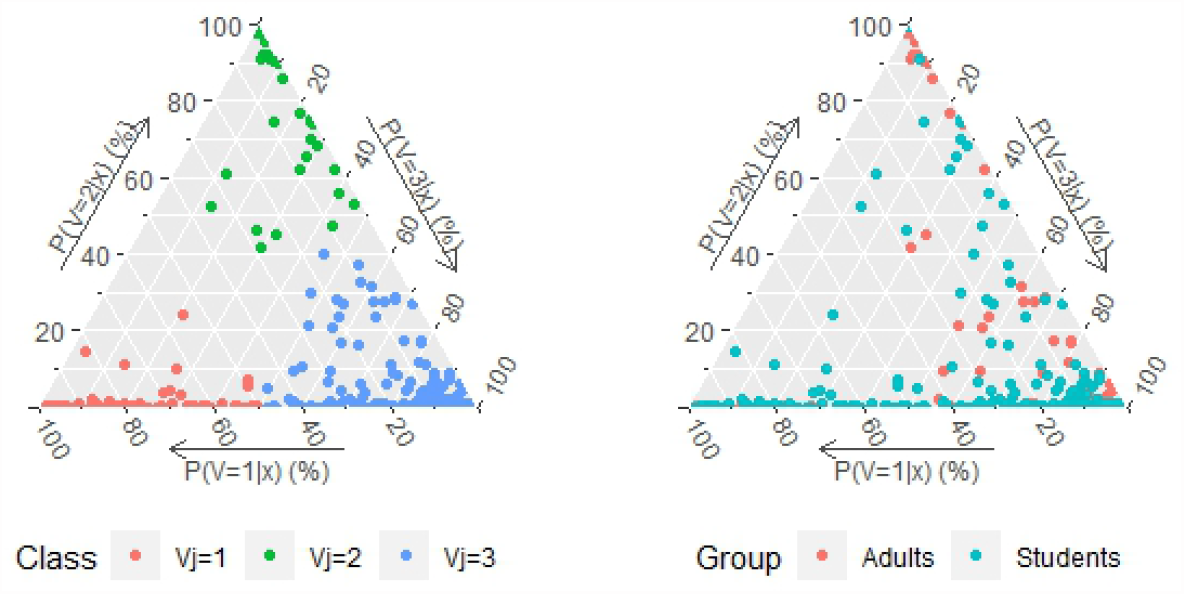
Ternary plots of the posterior probability (per cent) of belonging to each latent class for each of the individuals. Left plot: the modal assignment that results from assigning each individual to the latent class with the highest posterior membership probability is labeled in the graph. Right plot: the group (young students and healthy older adults) is labeled in the graph.

## Discussion

There is extensive literature which generally points to a detrimental effect of aging on pseudo-random productions [16–18]. In this work, we focused on analyzing the mental model of randomness that people implicitly use when producing random series. Our results revealed that there are significant age-related differences in the way individuals produce random-like sequences. A Markov chain of a given memory-order is often used to describe human-generated series. On this basis, individuals will deviate from randomness due to the influence of their previous choices, which they might recall at least vaguely. Indeed, this degree of dependence on previous responses could be closely related to the storage capacity of working memory [29, 30]. The *negative recency*, i.e., the tendency to alternate more than expected by chance, has received much attention and has usually been tested using a first-order Markov model [27].

Firstly, we fitted high-order Markov chains separately for the sample of young students and the sample of healthy older adults. Our results indicated that the length of the sliding window in which previous responses are taken into account to end up influencing the current head-tail choice is different between the two groups of subjects. Young students, in their random generation task, are generally able to recall in detail the sequence of their three immediately preceding choices, but only vaguely recall a summary of their longer-term responses up to memory-order 6. For the group of healthy older adults, they only accurately recall their immediate choice and more vaguely their most distant responses up to memory-order 3. Furthermore, our results obtained for both groups confirmed the hypothesis of a *negative recency* effect reported in the extensive literature on this topic.

Secondly, we proposed a novel approach that also takes into account intra-group heterogeneity, as it seems unrealistic to assume that within-group subjects generate sequences all of the same memory-order [39]. We hypothesized that there could be 3 types of individuals according to their way of mentally generating random sequences: individuals ‘without memory’ who would be able to replicate a series of fair coin tosses (latent class 1), individuals of short-time memory (latent class 2) and of long-time memory (latent class 3). Our results reveal an alternation bias in the sequences generated by respondents from classes 2 and 3, as well as a tendency to balance the distribution of their replies in short sequence segments. However, in this attempt to evaluate randomness in fragments of short sequences, in terms of memory requirements, the mental mechanism implied in the underlying process associated with class 3 is somewhat more sophisticated than that of class 2. We found that the probability of belonging to class 2 was significantly higher in the group of healthy older adults compared to the group of young students. This finding may be related to a previously published study [18] in which age was found to be the main factor affecting the complexity of random-like sequences, peaking at 25 years of age and declining more rapidly after age 60.

Finally, we observed a cluster of subjects with high posterior probabilities of belonging to class 1 (Fig 2). As mentioned above, subjects generating head-to-tail series according to the underlying process associated with class 1 would be able to reliably replicate a fair coin toss. However, we did not found sufficient statistical evidence to conclude that the probability of belonging to class 1 was significantly higher in the group of young students than in that of healthy older adults. On this point, further analysis would be necessary, such as analyzing how the results would change in dual-task experiments. Dual-task experiments require subjects to monitor or pay attention to more than one task at the same time. In general, published studies points to a detriment of randomness in random generation tasks when a second task is incorporated in a way that interferes with the main task [2, 20]. An analysis of performance in the random generation under dual-task conditions could provide more information on what leads some subjects to perform better than others. In addition, it would be interesting to analyze whether in these same dual-task conditions the differences between the two age groups increase or, on the contrary, remain the same. This is an interesting issue to investigate in future work.

## Supporting information

**S1 Table.** Estimated parameters (group of young students) for the latent class model of long-memory order *u* = 6.

| Parameter | Class 1 (c=1) | Class 2 (c=2) | Class 3 (c=3) |
| --- | --- | --- | --- |
| $P_{st}(V_j = c)$ | 0.338 (0.061) | 0.085 (0.027) | 0.577 (0.070) |
| $b_{01,st}$ | -0.040 (0.035) | | |
| $b_{02,st}$ | | 1.861 (0.066) | |
| $b_{12,st}$ | | -1.112 (0.127) | |
| $b_{22,st}$ | | -1.272 (0.108) | |
| $b_{03,st}$ | | | 1.724 (0.156) |
| $b_{13,st}$ | | | -0.829 (0.090) |
| $b_{23,st}$ | | | -0.538 (0.050) |

The first six observations for each subject are removed from the sample. Standard errors for the estimates in parentheses.

**S2 Table.** Estimated parameters (group of healthy older adults) for the latent class model of long-memory order *u* = 6.

| Parameter | Class 1 (c=1) | Class 2 (c=2) | Class 3 (c=3) |
| --- | --- | --- | --- |
| $P_{ad}(V_j = c)$ | 0.329 (0.087) | 0.261 (0.067) | 0.410 (0.098) |
| $b_{01,ad}$ | 0.009 (0.066) | | |
| $b_{02,ad}$ | | 2.573 (0.293) | |
| $b_{12,ad}$ | | -1.210 (0.213) | |
| $b_{22,ad}$ | | -1.803 (0.191) | |
| $b_{03,ad}$ | | | 1.766 (0.313) |
| $b_{13,ad}$ | | | -1.284 (0.186) |
| $b_{23,ad}$ | | | -0.512 (0.109) |

The first six observations for each subject are removed from the sample. Standard errors for the estimates in parentheses.

## Acknowledgments

This work was partially funded by the grant PID2022-137414OB-I00 from the Spanish Ministry of Science, Innovation and Universities and by the Spanish State Research Agency, through the Severo Ochoa and María de Maeztu Program for Centers and Units of Excellence in R&D (CEX2020–001084-M). The authors thank professor Anabel Blasco-Moreno (Department of Mathematics, Autonomous University of Barcelona) for additional suggestions and helpful comments.

## References

1. Sexton NJ, Cooper RP. An architecturally constrained model of random number generation and its application to modeling the effect of generation rate. Frontiers in Psychology. 2014;5:1–14.

2. Cooper RP. Executive functions and the generation of “random” sequential responses: A computational account. Journal of Mathematical Psychology. 2016;73:153–168.

3. Biesaga M, Talaga S, Nowak A. The effect of context and individual differences in human-generated randomness. Cognitive Science. 2021;45(12):e13072.

4. Baddeley A. Working memory. Oxford: Oxford University Press. 1986.

5. Baddeley A. Exploring the central executive. The Quarterly Journal of Experimental Psychology Section A. 1996;49(1):5–28.

6. Baddeley A, Emslie H, Kolodny J, Duncan J. Random generation and the executive control of working memory. The Quarterly Journal of Experimental Psychology Section A. 1998;51(4):819–852.

7. Towse J. On random generation and the central executive of working memory. The British Journal of Psychology. 1998;89(1):77–101.

8. Miyake A, Friedman NP, Emerson MJ, Witzki AH, Howerter A, Wager TD. The unity and diversity of executive functions and their contributions to complex “Frontal Lobe” tasks: a latent variable analysis. Cogn Psychol. 2000;41(1):49–100.

9. Salamé P, Danion JM. Inhibition of inappropriate responses is preserved in the think-no-think and impaired in the random number generation tasks in schizophrenia. J Int Neuropsychol Soc. 2007;13(2):277–287.

10. Koike S, Takizawa R, Nishimura Y, Marumo K, Kinou M, Kawakubo Y, Rogers MA, Kasai K. Association between severe dorsolateral prefrontal dysfunction during random number generation and earlier onset in schizophrenia. Clinical Neurophysiology. 2011;122(8):1533–1540.

11. Gilbert SJ, Bird G, Brindley R, Frith CD, Burgess PW. Atypical recruitment of medial prefrontal cortex in autism spectrum disorders: An fMRI study of two executive function tasks. Neuropsychologia. 2008;46(9):2281–2291.

12. Watkins E, Brown RG. Rumination and executive function in depression: an experimental study. Journal of Neurology, Neurosurgery & Psychiatry. 2002;72(3):400–402.

13. Brown RG, Soliveri P, Jahanshahi M. Executive processes in Parkinson’s disease–random number generation and response suppression. Neuropsychologia. 1998;36(12):1355–1362.

14. Williams IA, Wilkinson L, Limousin P, Jahanshahi M. Load-dependent interference of deep brain stimulation of the subthalamic nucleus with switching from automatic to controlled processing during random number generation in Parkinson’s disease. Journal of Parkinson’s disease. 2015;5(2):321–331.

15. Brugger P, Monsch AU, Salmon DP, Butters N. Random number generation in dementia of the Alzheimer type: a test of frontal executive functions. Neuropsychologia. 1996;34(2):97–103.

16. Van der Linden M, Beerten A, Pesenti M. Age-related differences in random generation. Brain and Cognition. 1998;38(1):1–16.

17. Heuer H, Janczyk M, Kunde W. Random noun generation in younger and older adults. Quarterly Journal of Experimental Psychology. 2010;63(3):465–478.

18. Gauvrit N, Zenil H, Soler-Toscano F, Delahaye JP, Brugger P. Human behavioral complexity peaks at age 25. PLoS Comput Biol. 2017;13(4):e1005408.

19. Towse JN, Neil D. Analyzing human random generation behavior: A review of methods used and a computer program for describing performance. Behavior Research Methods, Instruments and Computers. 1998;30(4):583–591.

20. Towse JN, Valentine JD. Random generation of numbers: A search for underlying processes. European Journal of Cognitive Psychology. 1997;9(4):381–400.

21. Oomens W, Maes JHR, Hasselman F, Egger JIM. RandseqR: An R package for describing performance on the random number generation task. Frontiers in Psychology. 2021;12:629012.

22. Gauvrit N, Zenil H, Delahaye JP, Soler-Toscano F. Algorithmic complexity for short binary strings applied to psychology: a primer. Behavior research methods. 2014;46(3):732–744.

23. Soler-Toscano F, Zenil H, Delahaye JP, Gauvrit N. Calculating Kolmogorov complexity from the output frequency distributions of small Turing machines. PloS one. 2014;9(5):e96223.

24. Gauvrit N, Singmann H, Soler-Toscano F, Zenil H. Algorithmic complexity for psychology: a user-friendly implementation of the coding theorem method. Behavior research methods. 2016;48(1):314–329.

25. Raftery A. A model for high-order Markov chains. Journal of the Royal Statistical Society: Series B. 1985;47(3):528–539.

26. Baena-Mirabete S, Espinal A, Puig P. Exploring the randomness of mentally generated head–tail sequences. Statistical Modelling. 2020;20(3):225–248.

27. Budescu DV. A Markov model for generation of random binary sequences. Journal of Experimental Psychology: Human Perception and Performance. 1987;13(1):25–39.

28. Schulz M-A, Schmalbach B, Brugger P, Witt K. Analysing humanly generated random number sequences: a pattern-based approach. PLoS One. 2012;7(7):e41531.

29. Miller GA. The magical number seven, plus or minus two: Some limits on our capacity for processing information. Psychological Review. 1956;63(2):81–97.

30. Saaty TL, Ozdemir MS. Why the magic number seven plus or minus two. Mathematical and Computer Modelling. 2003;38(3–4):233–244.

31. Meyniel F, Maheu M, Dehaene S. Human inferences about sequences: a minimal transition probability model. PLoS Comput Biol. 2016;12(12):e1005260.

32. De la Torre J, Douglas JA. Higher-order latent trait models for cognitive diagnosis. Psychometrika. 2004;69(3):333–353.

33. Swanson HL, Arizmendi GD, Li J-T. The stability of learning disabilities among emergent bilingual children: A latent transition analysis. Journal of Educational Psychology. 2021;113(6):1244–1268.

34. Spann MN, Silberman A, Feldman J, Korzeniewski SJ, Turner JB, Whitaker AH. Executive and non-executive functions in low birthweight/preterm adolescents with differing temporal patterns of inattention. PLoS One. 2020;15(4):e0231648.

35. Cugnata F, Martoni RM, Ferrario M, Di Serio C, Brombin C. Modeling physiological responses induced by an emotion recognition task using latent class mixed models. PLoS One. 2018;13(11):e0207123.

36. Benassi M, Garofalo S, Ambrosini F, Sant’Angelo RP, Raggini R, De Paoli G, Ravani C, Giovagnoli S, Orsoni M, Piraccini G. Using two-step cluster analysis and latent class cluster analysis to classify the cognitive heterogeneity of cross-diagnostic psychiatric inpatients. Frontiers in Psychology. 2020;11:1085.

37. Satoh M. Clustering of health behaviors among Japanese adults and their association with socio-demographics and happiness. PLoS One. 2022;17(4):e0266009.

38. Dempster AP, Laird NM, Rubin DB. Maximum likelihood from incomplete data via the EM algorithm. Journal of the Royal Statistical Society: Series B. 1977;39(1):1–38.

39. Shteingart H, Loewenstein Y. Heterogeneous suppression of sequential effects in random sequence generation, but not in operant learning. PLoS One. 2016;11(8):e0157643.

